# Amyloid-β deposition in basal frontotemporal cortex is associated with selective disruption of temporal mnemonic discrimination

**DOI:** 10.1101/2024.08.23.609449

**Authors:** Casey R. Vanderlip, Lisa Taylor, Soyun Kim, Alyssa L. Harris, Nandita Tuteja, Novelle Meza, Yuritza Y. Escalante, Liv McMillan, Michael A. Yassa, Jenna N. Adams

## Abstract

Cerebral amyloid-beta (Aβ) accumulation, a hallmark pathology of Alzheimer’s disease (AD), precedes clinical impairment by two to three decades. However, it is unclear whether Aβ contributes to subtle memory deficits observed during the preclinical stage. The heterogenous emergence of Aβ deposition may selectively impact certain memory domains, which rely on distinct underlying neural circuits. In this context, we tested whether specific domains of mnemonic discrimination, a neural computation essential for episodic memory, exhibit specific deficits related to early Aβ deposition. We tested 108 cognitively unimpaired human older adults (66% female) who underwent 18F-florbetapir positron emission tomography (Aβ-PET), and a control group of 35 young adults, on a suite of mnemonic discrimination tasks taxing object, spatial, and temporal domains. We hypothesized that Aβ pathology would be selectively associated with temporal discrimination performance due to Aβ’s propensity to accumulate in the basal frontotemporal cortex, which supports temporal processing. Consistent with this hypothesis, we found a dissociation in which generalized age-related deficits were found for object and spatial mnemonic discrimination, while Aβ-PET levels were selectively associated with deficits in temporal mnemonic discrimination. Further, we found that higher Aβ-PET levels in medial orbitofrontal and inferior temporal cortex, regions supporting temporal processing, were associated with greater temporal mnemonic discrimination deficits, pointing to the selective vulnerability of circuits related to temporal processing early in AD progression. These results suggest that Aβ accumulation within basal frontotemporal regions may disrupt temporal mnemonic discrimination in preclinical AD, and may serve as a sensitive behavioral biomarker of emerging AD progression.

## Introduction

In Alzheimer’s disease (AD), amyloid-beta (Aβ) pathology begins to accumulate within the posteromedial and basal frontotemporal cortex (Braak & Braak, 1991; Thal et al., 2002) decades prior to clinical impairment (Jack et al., 2018). Given the early role of Aβ in AD pathogenesis, identifying subtle cognitive deficits that reflect emerging Aβ accumulation could enable early detection of AD and sensitively assess clinical intervention outcomes for Aβ-lowering therapeutics.

Previous research has demonstrated small and inconsistent associations between Aβ pathology and concurrent cognitive ability during preclinical AD (Adams et al., 2023; Baker et al., 2017; Chételat et al., 2013; Hedden et al., 2013), inspiring debate on whether Aβ impacts cognition during the preclinical stage compared to the larger effects observed for tau or neurodegeneration (Maass et al., 2019; Mattsson-Carlgren et al., 2023). However, most previous studies have used dichotomous or global measures of Aβ (Donohue et al., 2014, 2017; Insel et al., 2020; Landau et al., 2012), which may miss the contribution of heterogeneous Aβ accumulation patterns. Further, many studies have used composite memory measures from neuropsychological batteries (Donohue et al., 2014) which may preclude identifying subtle impacts of Aβ on episodic memory deficits. Therefore, a regionally-based investigation of Aβ pathology’s effects on specific mechanisms underlying episodic memory is needed to elucidate its potential role in early, subtle memory deficits.

One critical aspect of episodic memory is pattern separation, a neural computation performed in the hippocampus which reduces interference among similar representations to enable discrimination of overlapping experiences (Bakker et al., 2008; Berron et al., 2016; Yassa & Stark, 2011). Pattern separation occurs across various domains, including object stimuli (Yassa et al., 2010), spatial relationships (Reagh & Yassa, 2014), and temporal ordering (Roberts et al., 2014). While the hippocampus is crucial for pattern separation across domains, cortical regions outside the hippocampus preferentially process specific stimuli and relay this domain-specific information to the hippocampus (Ranganath & Ritchey, 2012).

Extensive research has investigated how aging and AD affect object and spatial mnemonic discrimination and associated neural circuits (Adams et al., 2022; Maass et al., 2019; Reagh et al., 2018; Stark et al., 2010). Older adults demonstrate mnemonic discrimination for object prior to spatial stimuli (Güsten et al., 2021; Holden et al., 2012; Reagh et al., 2016, 2018). Anterior-temporal cortical systems supporting object processing and posterior-medial cortical systems supporting spatial processing are differently vulnerable to tau and Aβ pathology, respectively (Maass et al., 2019; Ranganath & Ritchey, 2012). While Aβ’s propensity to deposit within posteromedial cortex has motivated work investigating Aβ-related spatial mnemonic discrimination deficits, evidence for a direct association between Aβ and spatial mnemonic discrimination performance in older adults is weak (Adams et al., 2022; Webb et al., 2020), suggesting it may not be a sensitive marker of Aβ pathology in the preclinical stage.

Conversely, temporal processing engages basal frontotemporal regions (Duarte et al., 2010; E. L. Johnson et al., 2022) that are also highly susceptible to early Aβ accumulation (Braak & Braak, 1991; Grothe et al., 2017), yet temporal mnemonic discrimination has not been studied in the context of emerging AD. Neurons in medial orbitofrontal cortex (mOFC) track sequences of events (Jafarpour et al., 2023), and lesions in mOFC cause impairments in temporal memory even when object and spatial memory remain unaffected (Shimamura et al., 1990). Further, the inferior temporal (IT) cortex integrates items within a temporal framework (Kulkarni & Lega, 2023; Naya et al., 2017; Naya & Suzuki, 2011), without which temporal discrimination within the hippocampus would be impaired.

The involvement of basal frontotemporal regions in temporal processing and their vulnerability to early Aβ deposition suggest that temporal mnemonic discrimination performance could reflect underlying Aβ pathology. Here, we tested whether Aβ deposition contributes to deficits in temporal mnemonic discrimination in cognitively unimpaired older adults. We hypothesized that Aβ pathology would be selectively associated with temporal mnemonic discrimination performance, rather than with object or spatial, particularly when measured in basal frontotemporal regions supporting temporal processing.

## Materials and Methods

### Participants

35 young adults (18 to 35 years of age; mean 23.34 ± 4.42 years old; 71.4% female) and 108 cognitively unimpaired older adults (over 60 years of age; mean 71.22 ± 6.25 years old; 65.7% female) from the Biomarker Exploration in Aging, Cognition, and Neurodegeneration (BEACoN) study at the University of California, Irvine with assessment of mnemonic discrimination performance were included in this analysis (**Table 1**). Older adults additionally received magnetic resonance imaging (MRI) and positron emission tomography (PET) to assess Aβ pathology. All older adults were within their age-adjusted normal range in cognitive function based on neuropsychological testing. Further, all participants had no major health issues, and no history of significant neurological or psychiatric disorders. All participants provided written informed consent in accordance with the University of California, Irvine Institutional Review Board.

**Table 1.** Demographic information for study participants.

| | YA | Aβ- OA | Aβ+ OA |
| --- | --- | --- | --- |
| N | 35 | 70 | 38 |
| Age (years) M (SD) | 23.34 (4.42) | 70.29 (5.75) | 72.20 (6.86) |
| Education (Years) M (SD) | 14.46 (2.70) | 16.57 (2.41) | 15.21 (2.22) |
| Female N (%) | 25 (71.42%) | 42 (60.00%) | 29 (76.32%) |
| Race |  |  |  |
| White N (%) | 26.47% | 51 (72.86%) | 35 (92.11%) |
| Asian N (%) | 52.94% | 14 (20.00%) | 3 (7.89%) |
| Black N (%) | 2.94% | 1 (1.43%) | n/a |
| More than one N (%) | 5.88% | 2 (2.86%) | n/a |
| Other | 11.76% | 2 (2.86%) | n/a |
| MMSE M (SD) | n/a | 28.41 (1.35) | 28.62 (1.83) |

Aβ - amyloid- $\beta$ , YA - Younger adults, OA - Older adults, MMSE - Mini-Mental State Examination, M - mean, SD - standard deviation

### Mnemonic Discrimination Tasks

Young and older adult participants performed a suite of Mnemonic Discrimination Tasks (MDT), with distinct assessments of object, spatial, and temporal mnemonic discrimination.

### Object and Spatial

The object and spatial MDT (**Figure 1A-B**) have been extensively described in previous research (Adams et al., 2022; Kim et al., 2023; Reagh et al., 2018). Briefly, both the object and spatial domain MDT included an encoding phase where participants viewed a series of 120 images, making indoor/outdoor judgments for each image (presented for 2 s; interstimulus interval, 0.5 s). Participants had up to two seconds to respond after the stimulus was removed from the screen. In the object domain test phase, participants viewed a series of images classified as repetitions (identical to those seen during encoding), foils (completely new), or lures (similar but not identical to an image from encoding). These lures are particularly critical as they are known to require hippocampal pattern separation. In the object task, participants were asked to make “old”/“new” judgments based on whether they had seen the images during the encoding phase. Conversely, in the spatial domain test phase, participants viewed the same images as in the encoding phase but had to determine if the image’s location on the screen had changed (“same”/“different” judgment). The images were either in the same location (repetition) or had moved slightly (within the same screen quadrant) or significantly (to a different quadrant). This setup was designed to tax spatial pattern separation capacity. For both the object and spatial versions of the MDT, we calculated the response bias-corrected Lure Discrimination Index (LDI), quantified as p(“New or Different” | Lure) – p(“New or Different” | Target). To remove individuals with contaminant behavior, we excluded subjects (object: n = 5, spatial: n = 2) who responded greater than 2 standard deviations faster than the group average.

**Figure 1.**
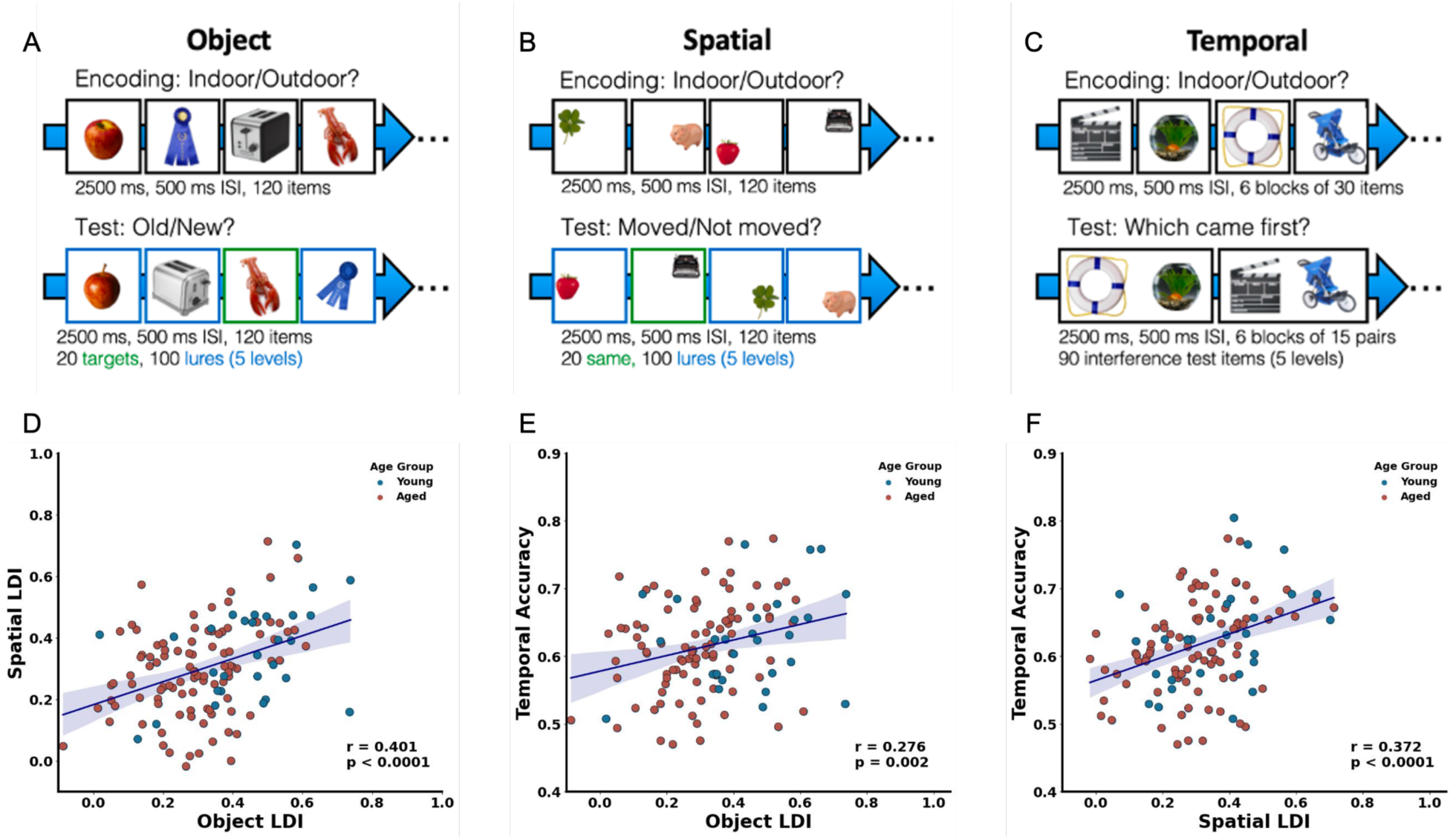
Mnemonic discrimination performance is positively correlated across domains. Schematic of **(A)** Object, **(B)** Spatial, and **(C)** Temporal mnemonic discrimination tasks. **(D)** Object and Spatial mnemonic discrimination are positively correlated. **(E)** Significant positive association between object mnemonic discrimination and temporal discrimination. **(F)** Strong positive association between spatial mnemonic discrimination and temporal discrimination. For **D-F**, Young adults are in blue and older adults are in red. Linear regression models were used to determine 95% confidence intervals shaded in gray.

### Temporal

The temporal MDT is designed to assess temporal mnemonic discrimination abilities (**Figure 1C**). The task was divided into 6 blocks, each comprising an encoding phase followed by a test phase. During the encoding phase, participants viewed a series of 30 images, making indoor/outdoor judgments for each (presented for 2 s; interstimulus interval, 0.5 s). Subsequently, in the test phase, they were presented with two images previously seen during encoding side by side, and asked to determine which image had appeared first during the encoding phase by making a “left”/ “right” judgment. Each image was displayed only once during the encoding phase and once during the test phase for each block, ensuring no repetitions within the context of the task. The majority (75%) of test phase trials assessed temporal lags between stimuli ranging from zero to 16 images during encoding. Additionally, a subset of trials (25%) featured images from the first and last positions of the encoding phase, assessing lags of 27 to 29 (Primacy/Recency trials). However, due to the potential influence of attentional rather than purely mnemonic demands on these Primacy/Recency trials (Morrison et al., 2014), we excluded them from our analysis. Consequently, we calculated temporal discrimination performance based on the accuracy of trials with lags of 16 or fewer across all blocks for each participant. Similar to the analysis of the object and spatial MDT, we excluded subjects (n = 4) who responded greater than 2 standard deviations faster than the group average.

### Structural MRI Acquisition and Processing

Older adults underwent MRI at the Facility for Imaging Brain Research (FIBRE) at UC Irvine on a 3T Prisma scanner (Siemens Medical System, Munich, Germany) equipped with a 32- channel head coil. A whole brain, high resolution T1-weighted volumetric magnetization prepared rapid gradient echo images (MPRAGE) was acquired for structural analyses (voxel size = 0.8mm^3^ resolution, TR/TE/TI = 2300/2.38/902 ms, flip angle = 8°, 240 slices acquired sagittally).

T1-weighted images were processed through FreeSurfer v.6.0.2 to obtain cortical native-space regions of interest for FBP normalization and quantification. T1-weighted images were also processed using Statistical Parametric Mapping 12 (SPM12) software, in which they were segmented into gray, white, and CSF components and warped to MNI152 2mm standard space. All FreeSurfer segmentation images and resulting MNI-template space T1-weighted images were quality checked by trained researchers, and FreeSurfer segmentations were manually edited when necessary.

### PET Acquisition and Processing

To assess Aβ pathology in older adult participants, 18F-florbetapir (FBP) PET was conducted using an ECAT High-Resolution Research Tomograph (CTI/Siemens) at the University of California, Irvine. A dose of 10 millicuries of the tracer was administered, followed by the acquisition of four 5-minute frames between 50 and 70 minutes post-injection. FBP images underwent attenuation correction, scatter correction, and 2 mm^3^ Gaussian smoothing. FBP images were then processed using SPM12 software. FBP images were realigned and coregistered to each participant’s T1-weighted MRI, and normalized with a whole cerebellum reference region to generate standardized uptake value ratio (SUVR) images. Next, additional 6 mm^3^ Gausian smoothing was applied, resulting in an effective resolution of 8 mm^3^. The mean SUVR from a cortical composite region previously validated for assessing global Aβ (Landau et al., 2012) was calculated to quantify overall Aβ burden. Aβ+ status was determined based on a threshold of >1.11 SUVR (Landau et al., 2012).

In addition to global measures, mean SUVR values were calculated for three composite regions representing stages of early, intermediate, and late Aβ vulnerability, as identified in prior work (Mattsson et al., 2019). For early-stage Aβ accumulation, the composite region encompassed the precuneus, posterior cingulate, isthmus cingulate, insula, and both medial and lateral orbitofrontal cortices. The late-stage region was composed of the lingual, pericalcarine, paracentral, precentral, and postcentral cortices. Lastly, the intermediate stage region was defined as the remaining cortical areas. Furthermore, specific regional SUVRs for the mOFC, IT, posterior cingulate cortex, and a combined control region of the primary motor and sensory cortices were also quantified.

We also conducted voxelwise FBP-PET analyses to determine spatially specific relationships between temporal mnemonic discrimination performance and FBP SUVR. Native space FBP images were warped to 2mm MNI space with SPM12, using deformation parameters generated from warping the T1-weighted image to MNI. Voxelwise associations between temporal discrimination performance and FBP SUVR were performed in SPM12, using a linear regression approach. A voxelwise threshold of p_unc_ < 0.005 and a cluster extent of greater than 25 voxels was used to determine significant clusters.

### Experimental Design and Statistical Analyses

Data was analyzed using Python. Shapiro-Wilk tests for normality revealed that the data were normally distributed, and so parametric statistical analyses were performed. Associations between two variables were assessed using Pearson correlations and linear regressions. Two-way analysis of variance (ANOVA) tests were used to assess two factor interactions. One-way ANOVAs with Tukey’s HSD post-hoc tests were used to identify within-factor differences. Independent samples t-test were used to assess differences between two groups based on Aβ status. Proportional shared effects of FBP SUVR within Aβ stages were performed with commonality analysis (Lindenberger & Pötter, 1998) using the “yhat” package in R, which conducts commonality analyses based upon all-possible-subsets regression. P < 0.05 was considered significant.

## RESULTS

### Age-related impairments in object and spatial mnemonic discrimination

To first investigate associations between behavioral performance on the object, spatial, and temporal domains of mnemonic discrimination, we correlated performance across the three tasks in young and older adults. Consistent with previous work, we found a strong positive association between object and spatial mnemonic discrimination (r_p_ = 0.401, p < 0.0001; **Figure 1D**). Further, we observed significant positive correlations between temporal mnemonic discrimination with both object (r_p_ = 0.276, p = 0.002; **Figure 1E**) and spatial mnemonic discrimination (r_p_ = 0.372, p < 0.0001; **Figure 1F**). These correlations remained significant within just the older adult subsample (temporal vs object: r_p_ = 0.271, p = 0.009; temporal vs spatial: r_p_ = 0.35, p = 0.001; object vs spatial: r_p_ = 0.354, p = 0.0004). These findings align with prior work suggesting that hippocampal pattern separation is domain agnostic (Azab et al., 2014, Yassa & Stark 2011). Critically, however, while statistically correlated, there was variability in the strength of the correlations. This suggests that there may be extra-hippocampal neural mechanisms that are not domain agnostic and therefore likely contribute to differing pattern separation capacities across domains.

We next tested how aging affects performance across object, spatial, and temporal domains by comparing performance between young and older adult groups. Consistent with prior findings (Stark et al., 2013), we observed a large impairment in object mnemonic discrimination in older adults (t = 5.23, p < 0.00001, Cohen’s d = 1.06), and a smaller, yet significant impairment in spatial mnemonic discrimination (t = 2.32, p = 0.022, Cohen’s d = 0.472) compared to younger adults. In contrast, temporal discrimination capacity did not significantly differ across age groups (t = 1.39, p = 0.17, Cohen’s d = 0.290). This aligns with prior work suggesting that, compared to other domains, object mnemonic discrimination is highly vulnerable to the aging process (Reagh et al., 2018), and suggests that temporal discrimination performance does not solely reflect age-related processes.

### Temporal mnemonic discrimination ability is selectively associated with elevated Aβ levels

While all older adults within this sample were cognitively unimpaired, a subset of individuals (n=38, 35%) were classified as Aβ+ based on their global 18F-florbetapir uptake (see **Methods**). We hypothesized that Aβ deposition would selectively disrupt temporal mnemonic discrimination and therefore first assessed whether performance on the mnemonic discrimination tasks varied according to Aβ status. To control for age-related changes and standardize measures across tasks, we z-scored performance of all older adults to the performance of young adults within each domain.

We found no significant difference in performance between Aβ- and Aβ+ older adults on either object or spatial mnemonic discrimination (object: t(87) = 0.80, p = 0.43; spatial: t(84) = 0.34, p = 0.74; **Figure 2A-B**). In contrast, in the temporal domain, Aβ+ older adults exhibited poorer performance compared to their Aβ- counterparts (t(85) = 2.03, p = 0.047; **Figure 2C**). Further, an analysis of continuous global Aβ levels demonstrated a significant negative correlation between global FBP SUVR and temporal mnemonic discrimination (temporal: r_p_ = -0.217, p = 0.043; **Figure 2F**), while no such association was found for object or spatial discrimination (object: r_p_ = -0.079, p = 0.460, spatial: r_p_ = -0.048, p = 0.658; **Figure 2D-E**). These findings collectively indicate a susceptibility of temporal discrimination to Aβ pathology, supporting our hypothesis that it may serve as a sensitive marker for early Aβ deposition in the absence of clinical symptoms. Given that Aβ was selectively related to temporal performance, our subsequent analyses sought to more closely examine the relationship between temporal discrimination and regional patterns of Aβ deposition to determine the underlying basis of this relationship.

**Figure 2.**
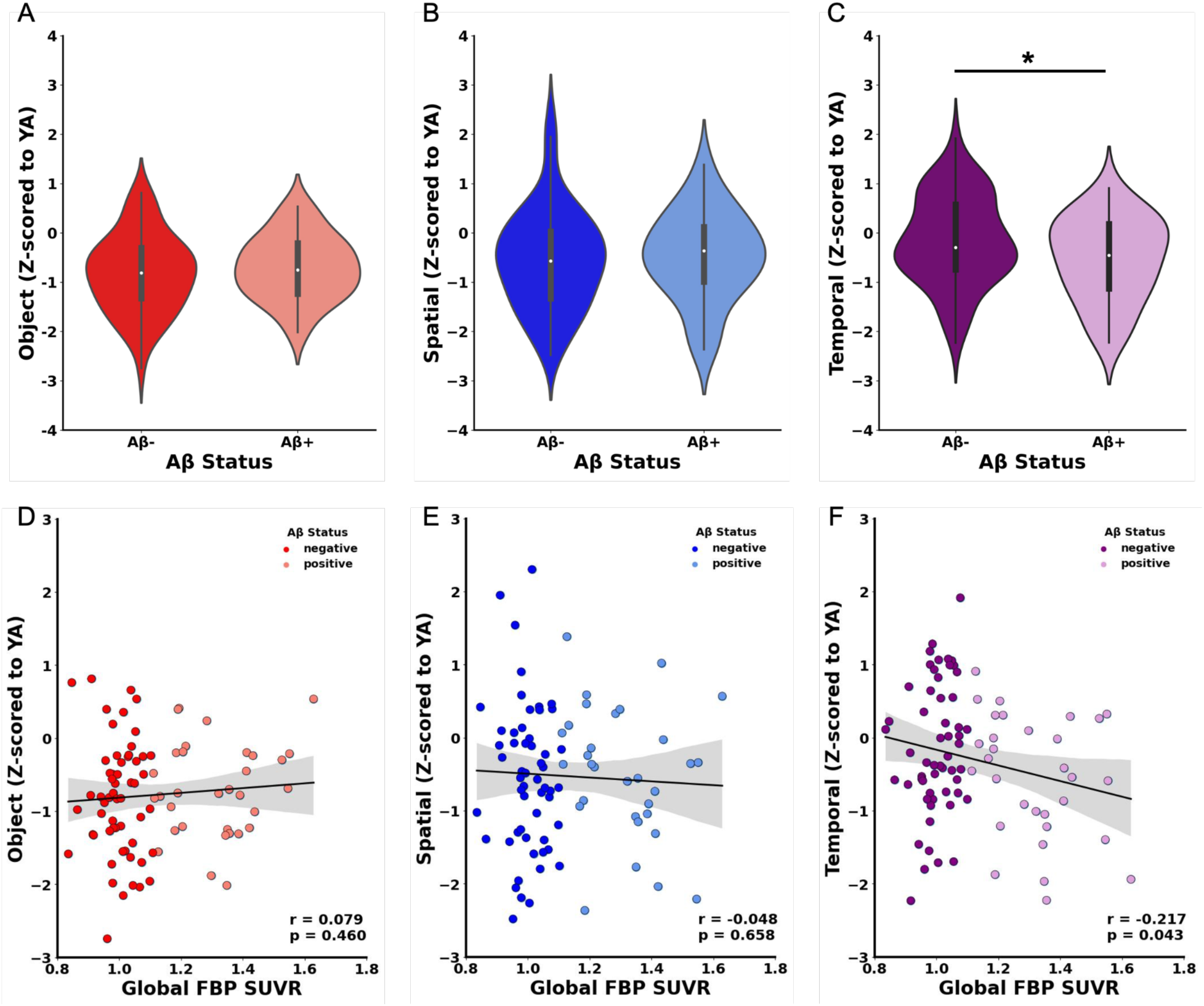
Global Aβ is selectively related to temporal discrimination performance in cognitively unimpaired older adults. Violin plots demonstrating that Aβ+ individuals performed similarly to Aβ- older adults on **(A)** object mnemonic discrimination and **(B)** spatial mnemonic discrimination. **(C)** Aβ+ older adults had impaired temporal discrimination compared to Aβ- individuals. There was no significant correlation between global SUVR and object **(D)** or spatial **(E)** mnemonic discrimination. **(F)** Increased global FBP SUVR was significantly associated with decreased temporal mnemonic discrimination. Linear regression models were used to determine 95% confidence intervals shaded in gray.

### Elevated Aβ levels in in regions impacted early in AD is associated with impaired temporal mnemonic discrimination

Given the overall association between temporal mnemonic discrimination and Aβ levels, we aimed to determine how early this Aβ effect may begin, hypothesizing Aβ levels in regions known to accumulate Aβ early would be related to performance. There are multiple proposed staging methods for identifying which areas are affected earliest by Aβ deposition, and there is not complete consensus on which regions should be included in each stage. Here we used one proposed staging scheme which categorized cortical regions into early, intermediate, and late stages based on the timing of Aβ accumulation as a testbed (Mattsson et al., 2019). We quantified mean FBP SUVR in early, intermediate, and late-stage regions (Mattsson et al., 2019; see **Methods**) and correlated these stage-specific FBP values with temporal discrimination accuracy. Our analyses revealed significant negative correlations between FBP SUVR with temporal mnemonic discrimination performance in both early (r_p_ = -0.240, p = 0.025; **Figure 3A**) and intermediate (r_p_ = -0.228, p = 0.034; **Figure 3B**) stages. However, Aβ levels in late-stage areas showed a marginal, but not significant, relationship with temporal discrimination (r_p_ = -0.170, p = 0.107; **Figure 3C**). To further investigate if Aβ in vulnerable regions are selectively related to temporal mnemonic discrimination, we conducted a commonality analysis to determine the unique versus shared variance of FBP SUVR within each stage on temporal mnemonic discrimination performance. This analysis demonstrated that the majority of variance (45.2%; coefficient = 0.035) of temporal mnemonic discrimination performance is explained by shared variance of early and intermediate stage Aβ. Shared variance across all three stages (21.4%; coefficient = 0.017) and unique variance of early (8.2%; coefficient = 0.006), intermediate (7.5%; coefficient = 0.006) and late stage FBP SUVR (25.1%; coefficient = 0.019) contributed to a lesser extent. These results suggest a notable link between increased Aβ deposition in areas vulnerable early in AD and poorer temporal discrimination. However, more importantly, this indicates that Aβ accumulation in specific brain regions, particularly those affected in the early and intermediate stages, may contribute to the deficits observed in temporal discrimination, rather than just global Aβ accumulation.

**Figure 3.**
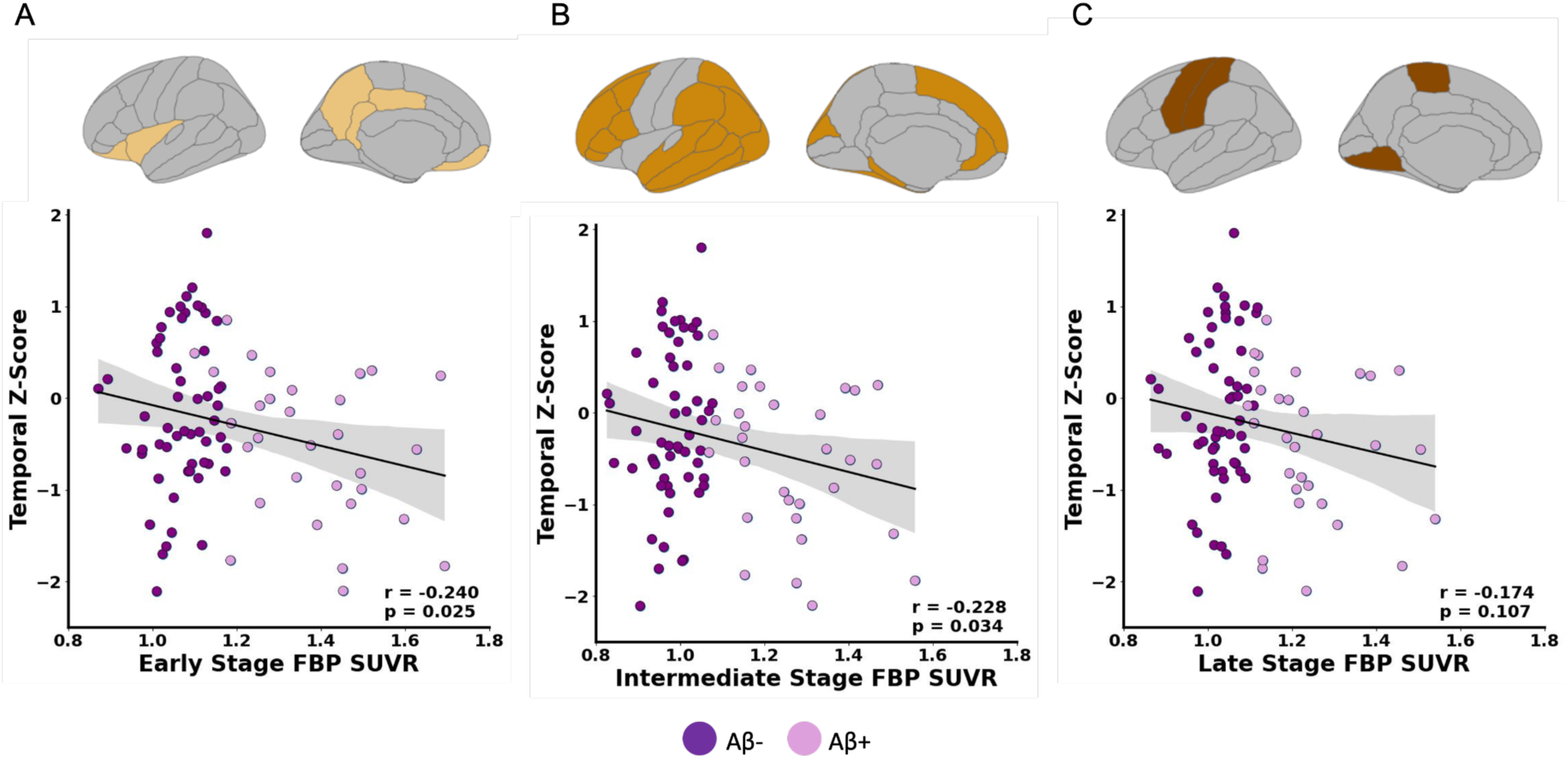
Aβ deposition in areas vulnerable to early accumulation is related to temporal discrimination. **A)** Significant negative association between FBP SUVR in early composite region composite and temporal mnemonic discrimination. **B)** Significant negative association between FBP SUVR in intermediate composite region and temporal mnemonic discrimination. **C)** No significant relationship between FBP SUVR in regions affected later in Aβ progression and temporal mnemonic discrimination.

### Aβ deposition in cortical regions supporting temporal memory are specifically associated with impairment in temporal discrimination

One plausible explanation for the link between Aβ deposition and impaired temporal discrimination is the involvement of specific brain circuits that are particularly susceptible to Aβ pathology. To test this hypothesis, we examined the relationship between temporal discrimination performance and Aβ levels in key cortical regions supporting temporal processing that are also known to accumulate Aβ (i.e. mOFC and IT). As control regions, we included the posterior cingulate cortex (PCC) as an early Aβ-accumulating posteromedial region that is not particularly involved in temporal processing, and primary sensory/motor cortex as a region that neither accumulates Aβ early nor contributes to temporal processing.

Consistent with this hypothesis, our analyses revealed a strong correlation between diminished temporal mnemonic discrimination with increased FBP SUVR in the mOFC (r_p_ = - 0.281, p = 0.008; **Figure 4A**) and IT (r_p_ = -0.325, p = 0.002; **Figure 4B**). In contrast, Aβ levels in the PCC, an area associated with Aβ deposition early in disease progression, but not temporal processing, was not significantly associated with temporal discrimination performance (r_p_ = - 0.176, p = 0.103; **Figure 4C**). Similarly, FBP SUVR in areas not traditionally associated with temporal processing or Aβ deposition, such as the primary motor and sensory cortices, showed no correlation with temporal mnemonic discrimination (r_p_ = -0.127, p = 0.243; **Figure 4D**). This lack of association in regions not implicated in temporal processing further supports the notion that Aβ-related impairments in temporal mnemonic discrimination are specifically linked to its deposition in brain areas integral to temporal processing and episodic memory.

**Figure 4.**
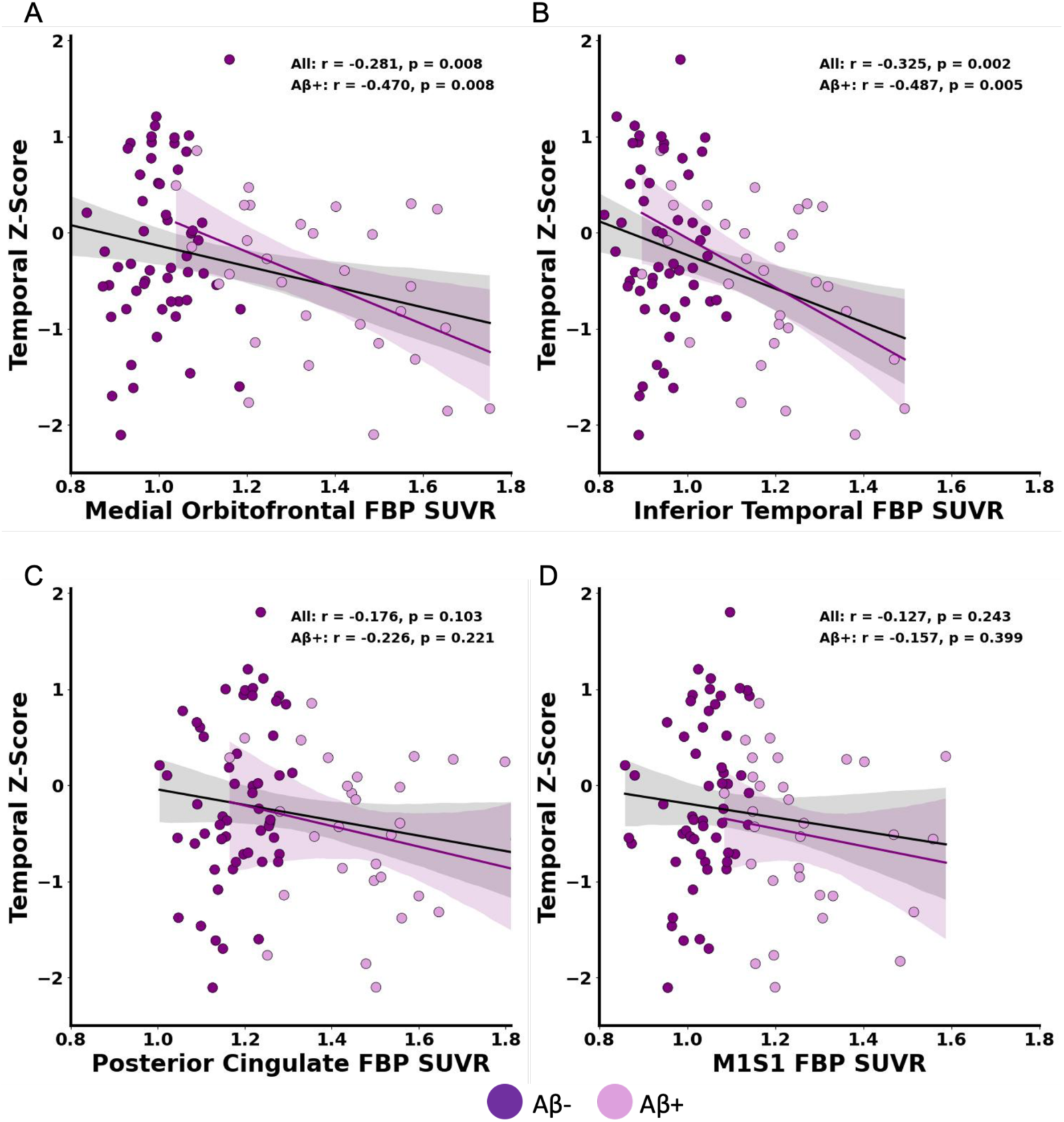
Selective relationship between Aβ deposition in regions related to temporal processing and temporal mnemonic discrimination performance. **(A)** Significant negative association between medial orbitofrontal cortex FBP SUVR and temporal mnemonic discrimination across all older adults (black line) and within the Aβ+ older adults (purple line). **(B)** Significant negative correlation between FBP SUVR in inferior temporal cortex and temporal discrimination across all older adults (black line) and within Aβ+ older adults (purple line). **(C)** Despite the posterior cingulate cortex being vulnerable to Aβ pathology, FBP SUVR did not significantly correlate with temporal mnemonic discrimination. **(D)** There was no relationship between FBP SUVR in primary motor/primary sensory cortex (M1S1) and temporal mnemonic discrimination.

Previous research indicates a complex relationship between Aβ deposition and cognitive deficits, suggesting that while Aβ may be associated with cognitive deficits, its levels do not consistently correlate with cognitive function beyond a certain threshold. Given this, we aimed to determine whether, within Aβ+ individuals, heightened regional Aβ levels were associated with worsening temporal discrimination deficits. We again found strong significant correlations between higher FBP SUVR in the mOFC and IT and deficits in temporal mnemonic discrimination when restricting analyses to Aβ+ individuals (mOFC: r_p_ = -0.470, p = 0.008, IT: r_p_ = -0.487, p = 0.005; **Figure 4A-B**). Conversely, we again observed no significant correlation between temporal mnemonic discrimination deficits and FBP SUVR in PCC (r_p_ = -0.226, p = 0.221; **Figure 4C**) and the primary motor and sensory cortices (r_p_ = -0.157, p = 0.399; **Figure 4D**) when restricting analyses to Aβ+ individuals. These results indicate that the degree of Aβ deposition in basal frontotemporal cortical regions is associated with worsening temporal mnemonic discrimination, suggesting that regional Aβ plays a direct role in the disruption of this cognitive function and is not just an incidental marker of preclinical AD.

To further validate the spatial specificity of this relationship, we employed a voxelwise approach to analyze the association between voxelwise FBP SUVR levels and temporal mnemonic discrimination performance. Our voxelwise analysis confirmed selective associations between worse temporal mnemonic discrimination performance with an increase in voxelwise FBP SUVR within clusters of the bilateral mOFC and bilateral IT (**Figure 5**; p_unc_<0.005, k>25), reinforcing our *a priori* hypothesis driven ROI-based results. Additional data-driven clusters included bilateral ventral visual cortex and fusiform cortex. Critically, this voxelwise association was again absent in regions known to be susceptible to Aβ deposition but not directly involved in temporal processing (i.e. posteromedial cortex regions such as PCC). This absence of association in areas unrelated to temporal processing further affirms the specificity of the link between Aβ pathology in regions supporting temporal processing, such as mOFC and IT, with temporal mnemonic discrimination deficits.

**Figure 5.**
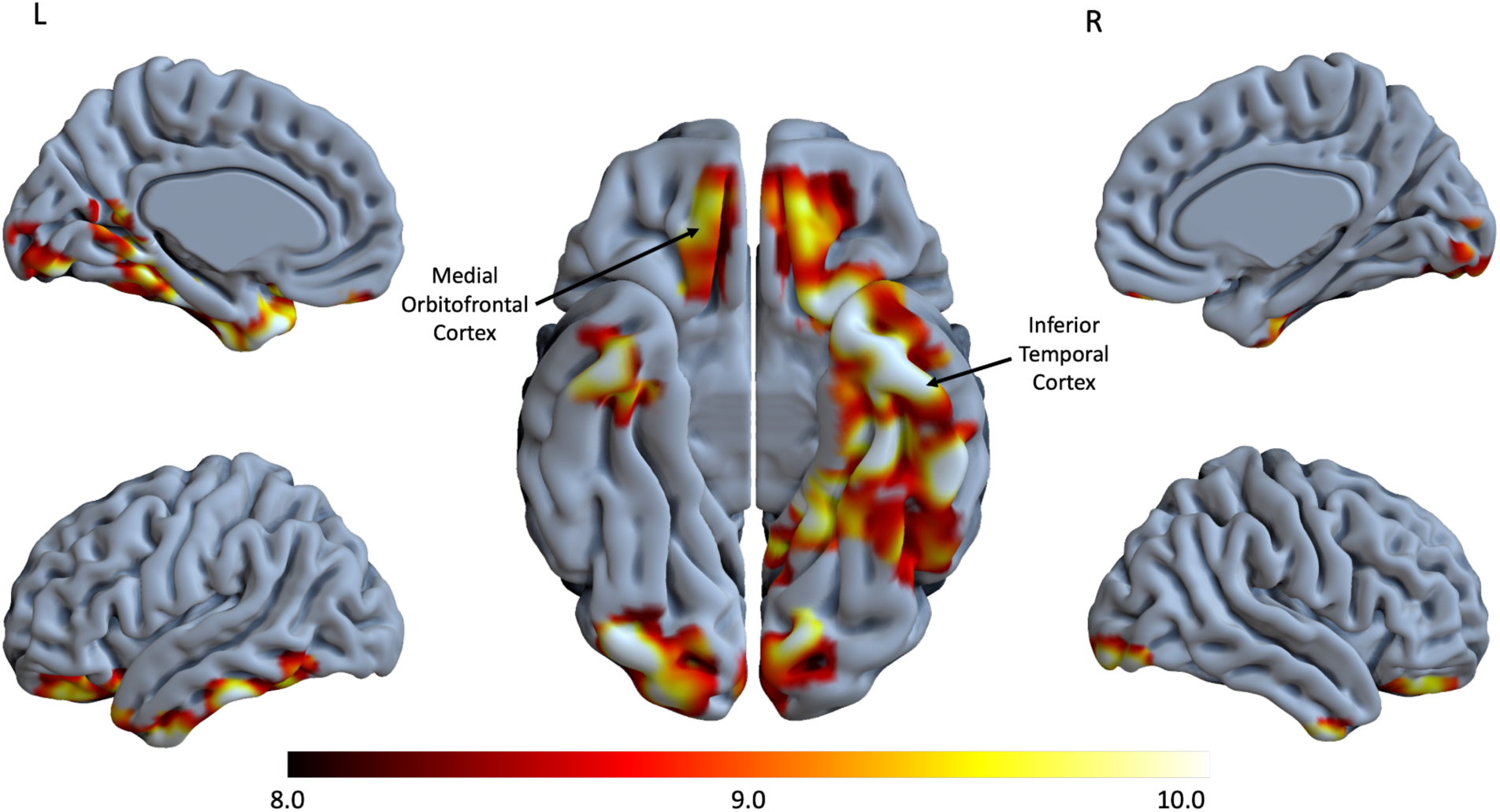
Voxelwise analysis of the association between FBP SUVR and temporal mnemonic discrimination. Medial and lateral view of both left and right hemispheres and inferior view of the whole brain. Increased FBP SUVR in basal frontotemporal regions such as medial orbitofrontal cortex and inferior temporal cortex is associated with temporal mnemonic discrimination deficits.

## Discussion

We demonstrate that Aβ deposition in the basal frontotemporal cortex is related to temporal mnemonic discrimination deficits in cognitively unimpaired older adults. We utilized distinct domains of mnemonic discrimination tasks to separately tax object, spatial, and temporal pattern separation in older adults who underwent Aβ-PET and young adult control participants. We found that older adults with elevated Aβ levels were selectively impaired on temporal, but not object or spatial, mnemonic discrimination. This temporal mnemonic discrimination deficit was driven by deposition of Aβ within cortical regions previously shown to support temporal processing such as mOFC and IT (Jafarpour et al., 2023; Naya et al., 2017), which are also regions proposed to have early vulnerability to Aβ deposition (Braak & Braak, 1991; Grothe et al., 2017). Our results suggest that temporal mnemonic discrimination deficits are a sensitive marker of early Aβ deposition in the preclinical stage of AD, and may be a useful domain to predict future disease progression and cognitive decline.

Previous investigations between Aβ and mnemonic discrimination have been restricted to the object or spatial domains (Adams et al., 2022; Berron et al., 2024; Jutten et al., 2022; Maass et al., 2019; Papp et al., 2020; Vanderlip et al., 2024). This work shows that while Aβ may have subtle relationships with object discrimination (Berron et al., 2024; Papp et al., 2020), especially in large studies powered to detect small effects (Papp et al., 2020; Young et al., 2023), these effects may be mediated through tau pathology in anterior-temporal cortical networks (Berron et al., 2019, 2024; Maass et al., 2019), suggesting an indirect relationship with Aβ. In the spatial domain, many studies have failed to observe a direct relationship between Aβ and performance (Adams et al., 2022; Berron et al., 2024; Kim et al., 2023), even though Aβ accumulates within posteromedial cortical regions that support spatial processing (Berron et al., 2018; Maass et al., 2019). However, it is important to note that one study which employed a mnemonic discrimination task varying both spatial location and object identity simultaneously did observe an association with global Aβ levels (Webb et al., 2020). Finally, studies using various features from mnemonic discrimination tasks to predict Aβ status have found features such as recognition memory or response biases had more predictive power than the actual mnemonic component of these tasks (Kim et al., 2023; Vanderlip et al., 2024), further supporting an indirect relationship with Aβ in object and spatial domains.

To our knowledge, temporal mnemonic discrimination has not previously been studied in the preclinical stage of AD. Our results substantiate previous work demonstrating robust age-group differences in object and spatial mnemonic discrimination (Leal & Yassa, 2018; Reagh et al., 2016; Stark et al., 2013, 2019). Critically, we provide a novel dissociation between temporal mnemonic discrimination, which we found to differ by Aβ status but not age, compared to object and spatial mnemonic discrimination, which we found to differ by age but not by Aβ status. However, previous work has shown age effects in temporal mnemonic discrimination at longer temporal lags, though Aβ biomarker information was unavailable (Roberts et al., 2014; Tolentino et al., 2012). Beyond Aβ positivity, our results extended to associations with continuous global FBP SUVR as well as FBP SUVR within early and intermediate stages of Aβ deposition (Mattsson et al., 2019), supporting the sensitivity of the temporal domain to Aβ pathology. Based on our current work and the previous literature, we propose that age-related changes in hippocampal integrity and medial temporal lobe tau may drive initial age-related deficits in object (and to a lesser extent, spatial) mnemonic discrimination (Berron et al., 2019, 2024; Stark & Stark, 2017). Conversely, temporal mnemonic discrimination may be disrupted in preclinical AD due to early Aβ deposition in basal frontotemporal cortex (Braak & Braak, 1991; Grothe et al., 2017; Thal et al., 2002), as these cortical regions support temporal processing. Finally, deficits in spatial mnemonic discrimination may emerge with symptomatic stages of mild cognitive impairment or AD, when failures of spatial processing are well established (Kim et al., 2023; Paleja & Spaniol, 2013).

Our primary mechanistic hypothesis was that Aβ accumulation affects temporal mnemonic discrimination because basal frontotemporal regions vulnerable to Aβ are also involved in temporal processing. Temporal mnemonic discrimination, which involves both the memory of sequences and items, as well as the capacity to bind items in sequence, depends on connections between the prefrontal cortex, hippocampus, and IT (Naya et al., 2017). Specifically, the mOFC contributes to temporal processing (Zhou et al., 2019) and has strong connections to IT, feeding input into the hippocampus through the perirhinal and entorhinal cortices (Heather Hsu et al., 2020; Huang et al., 2021; Saleem et al., 2008). IT is critical for item representation, with evidence underscoring its role in recognition memory (Easton & Gaffan, 2001; Miller et al., 1991; Naya & Suzuki, 2011). Both mOFC and IT have extensive connections to the medial temporal lobe, therefore we hypothesize that Aβ deposition could alter temporal mnemonic discrimination by degrading cortical input into the hippocampus, thus leading to weakened hippocampal representations. Indeed, we found strong associations between temporal mnemonic discrimination and Aβ in both regions. Importantly, these correlations persisted when assessing this association within only Aβ+ individuals. This critically supports that not only the presence, but the severity of Aβ, in these regions impacts temporal mnemonic discrimination in a dose-dependent manner.

To confirm the spatial specificity of associations between regional Aβ and temporal mnemonic discrimination, we conducted regional control and voxelwise analyses. We first contrasted the effect found in mOFC and IT with two control areas to provide a dissociation - one highly vulnerable to Aβ, but not directly involved in temporal processing (PCC) and one unaffected by Aβ and not involved in temporal processing (primary sensory/motor cortex). Consistent with our initial hypothesis, we did not observe a significant association between temporal mnemonic discrimination and Aβ levels in either region, which is notable given the high susceptibility of the PCC to Aβ accumulation (Leech & Sharp, 2014; Scheff et al., 2014). Remarkably, our voxelwise analyses also confirmed that Aβ within basal frontotemporal regions, but not posteromedial regions, was related to temporal mnemonic discrimination. The voxelwise spatial pattern of Aβ that was associated with temporal mnemonic discrimination strikingly resembles “Stage A” of the neuropathologically-driven Aβ accumulation model as described by Braak and colleagues (Braak & Braak, 1991), which similarly implicates medial frontal, basal temporal, and ventral visual cortex. Our voxelwise results are also consistent with the first stage of other PET-based staging studies of Aβ, which specifically include IT and orbitofrontal cortex (Cho et al., 2016; Grothe et al., 2017). This specificity highlights the importance of considering regional vulnerability and functional relevance when investigating the impacts of Aβ deposition on cognitive abilities, consistent with previous studies demonstrating stronger relationships between regional or voxelwise Aβ and cognition than global levels (Farrell et al., 2018; Sperling et al., 2019; Tideman et al., 2022).

A compelling future extension of this work is to explore the potential synergistic effects of tau pathology in the association between Aβ and temporal mnemonic discrimination (Ossenkoppele et al., 2022; Sperling et al., 2019). We suspect that tau deposition is also related to temporal mnemonic discrimination given previous work demonstrating anterolateral entorhinal cortex involvement in temporal processing (Montchal et al., 2019). Further, mOFC and IT are regions thought to reflect a transition between “age-related” medial temporal lobe tau and the widespread cortical tauopathy characteristic of AD (K. A. Johnson et al., 2016; Schöll et al., 2016), and regions concurrently expressing both pathologies may lead to greater deficits in neuronal processing. In the current study, we could not assess tau deposition due to a lack of tau-PET data, and therefore future work will investigate the important role tau may play in temporal mnemonic discrimination.

Limitations include the highly educated sample consisting of primarily non-Hispanic white individuals. Future work in more diverse populations will aid in the generalizability of these results. Further, it is possible that other mechanisms such as synaptic dysfunction or hyperactivity may mediate the relationship between Aβ and performance. Thus, future work needs to investigate other potential mechanisms that give rise to deficits in temporal mnemonic discrimination. Lastly, longitudinal data will be needed to directly assess whether temporal mnemonic discrimination deficits are predictive of future cognitive decline, as previous work demonstrates that longitudinal relationships between Aβ and cognitive decline may be stronger than cross-sectional effects (Donohue et al., 2017; Ossenkoppele et al., 2022; Sperling et al., 2019).

We identified a selective relationship between temporal mnemonic discrimination and Aβ pathology in cognitively unimpaired older adults. We demonstrate that this relationship is due to Aβ deposition in basal frontotemporal regions supporting temporal processing, which may preclude these regions from sending proper information to the hippocampus for temporal pattern separation computations. Our work suggests that deficits in temporal mnemonic discrimination may serve as an early marker for Aβ accumulation. In light of this, cognitive batteries could be expanded to include tasks taxing temporal discrimination to enhance early detection of individuals at high risk for cognitive decline and to better assess the efficacy of Aβ-lowering therapeutics.

## Author contributions

Conceptualization, C.R.V., J.N.A., & M.A.Y.; Data Acquisition, A.L.H., N.T., N.M., Y.Y.E., S.K.; Project administration, L.M.; Methodology & Software, C.R.V., L.T., J.N.A; Formal Analysis, C.R.V.; Writing – Original Draft, C.R.V. & J.N.A.; Writing – Reviewing and Editing, all authors; Funding Acquisition and Supervision, M.A.Y. & J.N.A.

## Acknowledgments

This research was supported by the National Institute on Aging R01AG053555 to M.A.Y. and F32AG074621 to J.N.A.

